# From de novo to ‘de nono’: most novel protein coding genes identified with phylostratigraphy represent old genes or recent duplicates

**DOI:** 10.1101/287193

**Authors:** Claudio Casola

**Affiliations:** Department of Ecosystem Science and Management, Texas A&M University, College Station, TX 77843-2138

## Abstract

The evolution of novel protein-coding genes from noncoding regions of the genome is one of the most compelling evidence for genetic innovations in nature. One popular approach to identify de novo genes is phylostratigraphy, which consists of determining the approximate time of origin (age) of a gene based on its distribution along a species phylogeny. Several studies have revealed significant flaws in determining the age of genes, including de novo genes, using phylostratigraphy alone. However, the rate of false positives in de novo gene surveys, based on phylostratigraphy, remains unknown. Here, I re-analyze the findings from three studies, two of which identified tens to hundreds of rodent-specific de novo genes adopting a phylostratigraphy-centered approach. Most of the putative de novo genes discovered in these investigations are no longer included in recently updated mouse gene sets. Using a combination of synteny information and sequence similarity searches, I show that about 60% of the remaining 381 putative de novo genes share homology with genes from other vertebrates, originated through gene duplication, and/or share no synteny information with non-rodent mammals. These results led to an estimated rate of ∼12 de novo genes per million year in mouse. Contrary to a previous study (Wilson et al. 2017), I found no evidence supporting the preadaptation hypothesis of de novo gene formation. Nearly half of the de novo genes confirmed in this study are within older genes, indicating that co-option of preexisting regulatory regions and a higher GC content may facilitate the origin of novel genes.

## Introduction

Protein-coding genes can emerge through mechanisms varying from gene duplication to horizontal transfer and the ‘domestication’ of transposable elements, all of which involve pre-existing coding regions. Conversely, the process of de novo gene formation consists of the evolution of novel coding sequences from previously noncoding regions, thus generating entirely novel proteins. The discovery of de novo genes is facilitated by extensive comparative genomic data of closely related species and their accurate gene annotation. Because of these requirements, de novo genes have been mainly characterized in model organisms such as *Saccharomyces cerevisiae* (Carvunis et al. 2012; Lu et al. 2017; Vakirlis et al. 2017), *Drosophila* (Begun et al. 2007; Reinhardt et al. 2013; Zhao et al. 2014) and mammals (Heinen et al. 2009; Knowles and McLysaght 2009; Li et al. 2010; Murphy and McLysaght 2012; Neme and Tautz 2013; Ruiz-Orera et al. 2015; Guerzoni and McLysaght 2016; Neme and Tautz 2016).

Even among model organisms, the identification of de novo genes remains challenging. One major caveat in de novo gene discovery is their association with signatures of biological function. Because de novo genes described thus far tend to be taxonomically restricted to a narrow set of species, their functionality is not obvious as for older genes that are conserved across multiple taxa. Transcription and translation are considered strong evidence of de novo genes functionality, although the detection of peptides encoded from putative coding sequences is not always an indication of protein activity (Xu and Zhang 2016). Given their limited taxonomic distribution, testing sequence conservation and selective regimes on de novo genes coding sequences is often unachievable. Population genomic data can provide evidence of purifying selection on de novo genes but are thus far limited to a relatively small number of species (Zhao et al. 2014; Chen et al. 2015).

An important signature of de novo gene evolution is the presence of *enabler* substitutions, which are nucleotide changes that alter a ‘proto-genic’ DNA region to facilitate its transcription and/or ability to encode for a protein (Vakirlis et al. 2017). Enabler substitutions can be recognized only by comparing de novo genes with their ancestral noncoding state, which can be identified by analyzing syntenic regions in the genome of closely related species that do not share such substitutions (Guerzoni and McLysaght 2016). It follows that putative de novo genes with no detectable synteny in other species have likely originated through other processes, including duplication of preexisting genes, horizontal gene transfer, or transposon insertion and subsequent domestication. Loss of synteny may also arise through deletion of orthologous genes in other species; however, this scenario appears improbable when comparing a large number of genomes.

A third fundamental hallmark of de novo genes is the lack of homology of their proteins with proteins from other organisms. This feature is often the first step in comparative genome-wide surveys aimed at identifying putative de novo genes in a given species or group of species. Similarly, de novo proteins should be devoid of functional domains that occur in older proteins.

Previous studies on de novo gene evolution show a range of complexity in the strategies used to characterize these genes. A common approach to assess the evolutionary age of genes in known as *phylostratigraphy* and it has often been applied to retrieve a set of candidate de novo genes. This method relies on homology searches, usually consisting of BLAST surveys of a species proteome, allowing the inference of a putative ‘age’ of each gene along the phylogeny of a group of species (Domazet-Loso et al. 2007). For example, mouse proteins that share homology with sequences in the rat proteome, but are absent in other mammals, would be categorized as ‘rodent-specific’. Notably, genes that appear to be lineage-specific may represent de novo genes, but could also be derived from any of the other processed mentioned above.

Phylostratigraphy is theoretically simple and effective, yet it is known to contain several methodological flaws (Elhaik et al. 2006; Moyers and Zhang 2015, 2016; Moyers and Zhang 2017). For instance, both rapid sequence evolution and short coding sequences lead to underestimating gene age (Moyers and Zhang 2015). This increases the likelihood for rapidly evolving genes to be erroneously recognized as novel species-specific genes when phylostratigraphy-only approaches are used. It is known that after gene duplication one or both of the two copies may experience accelerated sequence evolution, which may result in an underestimate of their age. In agreement with this observation, a recent study in primates has shown that genes that evolve faster also tend to duplicate more (O’Toole et al. 2018). Importantly, phylostratigraphic studies that ignore synteny data will be unable to provide evidence of enabler substitutions and are thus uninformative of the mechanism by which those genes evolved.

It is perhaps not surprising that researches chiefly based on phylostratigraphy have led to estimates of de novo gene formation rates that exceeds or are comparable to those of gene duplication. For instance, it has been suggested that the *S. cerevisiae* genome contains hundreds of de novo genes that emerged during the Ascomycota evolution and that at least 19 genes are *S. cerevisiae*-specific, compared to a handful of gene duplicates found only in *S. cerevisiae* (Carvunis et al. 2012). Similarly, a relatively recent study reported that 780 de novo genes emerged in mouse since its separation from the Brown Norway rat around 12 million years ago, at a rate of 65 genes/million year (Neme and Tautz 2013). According to these estimates, de novo genes represent about half of all young genes, the other half being formed by gene duplicates. Because the overall gene number did not appear to have increased significantly during mammal and yeast evolution, such a high pace of de novo gene formation must be accompanied by rampant levels of gene loss. If true, this would represent a ‘gene turnover paradox’, given that most genes are maintained across mammals. For instance, according to the Mouse Genome Database, 17,093/22,909 (∼75%) protein coding mouse genes share homology with human genes (Blake et al. 2017).

The ‘gene turnover paradox’ has been opposed by some authors. For example, analyses by Moyers and Zhang used simulations to show that gene age is underestimated in a significant proportion of cases based on phylostratigraphy (Moyers and Zhang 2015, 2016; Moyers and Zhang 2017). These authors pointed out that most putative *S. cerevisiae*-specific de novo genes overlap with older genes and show no signature of selection operating on their coding sequence (Moyers and Zhang 2016). These studies have proved critical to address major pitfalls of phylostratigraphy. However, the exact proportion of false positives in de novo gene studies remains unknown and it is unclear how many putative de novo genes should instead be considered fast evolving gene duplicates. A correct assessment of de novo genes is critical to establish their evolutionary history and more broadly to identify genomic features, if any, that may facilitate the emergence of novel genes.

Here, I address these issues by re-analyzing putative mouse de novo genes from three recent articles (Murphy and McLysaght 2012; Neme and Tautz 2013; Wilson et al. 2017) using a combination of sequence similarity searches and synteny information. I show that more than half of the 874 putative de novo genes previously described in mouse are absent in current versions of three major mouse gene annotation databases, an indication of how gene annotation volatility can affect de novo gene studies even among model organisms. Of the remaining putative de novo genes, only ∼40% could be validated. The dismissed putative de novo genes either shared homology with genes found in multiple non-rodent vertebrates, derived from duplication of pre-existing mouse genes, and/or lacked synteny information with non-rodent mammals. I collectively refer to the putative de novo genes that failed to pass the validation criteria as the ‘de nono’ genes. These findings also indicate that false positives in phylostratigraphy studies of de novo genes exceed previous estimates of type I error rates based on simulations in *S. cerevisiae* (Moyers and Zhang 2016). Contrary to what was suggested in a recent study (Wilson et al. 2017), I found no evidence of preadaptation in the validated mouse de novo genes. Instead, I observed that the trend reported by Wilson and collaborators, an inverse correlation between intrinsic structural disorder (ISD) of proteins and gene age suggestive of a lower tendency towards aggregation in proteins encoded by younger genes, is primarily due to high ISD levels in de novo genes whose coding region overlap exons of older genes.

## Results and Discussion

### Putative de novo gene annotation status

In this study, I re-analyzed <u>P</u>utative <u>D</u>e <u>N</u>ovo <u>G</u>enes (hereafter: PDNGs) from three articles focused on rodent genomes (Murphy and McLysaght 2012; Neme and Tautz 2013; Wilson et al. 2017). Hereafter, I will refer to these works as M2012 (Murphy and McLysaght 2012), N2013 (Neme and Tautz 2013) and W2017 (Wilson et al. 2017). I specifically focused on mouse-specific genes from the M2012 and N2013 studies, and on the rodent-specific genes from the W2017 study. A total of 491 previously reported rodent PDNGs, particularly those from the M2012 and N21023 studies, are not annotated as protein-coding genes in the updated versions of three major mouse gene annotation databases: GENCODE M16, RefSeq and UCSC Genome Browser ‘known’ genes (Table 1).

**Table 1.** Original PDNG sets, currently annotated PDNGs and de novo genes assessed in this study

|  | M2012 | N2013 | W2017 | Total |
| --- | --- | --- | --- | --- |
| Mouse PDNG | 69 | 773 | 84 | 874 |
| Annotated in mm10 | 9 | 331 | 72 | 381 |
| de novo genes | 3 (7) | 74 (139) | 13 (18) | 82 (152) |

Numbers in parenthesis refer to de novo genes remaining when genes from automatic annotation pipelines are included.

This is expected given that the three studies were based on less-well curated gene sets. Surprisingly, the N2013 and W2017 datasets contained several PDNGs lacking a start codon. These genes were excluded from further analysis in this work. The final count of PDNGs with confirmed annotation as protein-coding genes was 9, 331, and 72 from the M2012, N2013, and W2017 studies, respectively (Table 1). After excluding overlap between the three gene sets, I retrieved 381 PDNGs, which represent 44% of the 874 genes originally reported as de novo genes. Below, I describe the four criteria I used to assess the proportion of PDNGs that represent ‘de nono’ genes: presence of paralogous genes in mouse (inparalogs); homology with genes found in multiple non-rodent vertebrates; lack of synteny information with non-rodent mammals; presence of conserved domains found in non-rodent proteins (Figs. 1 2).

**Figure 1.**
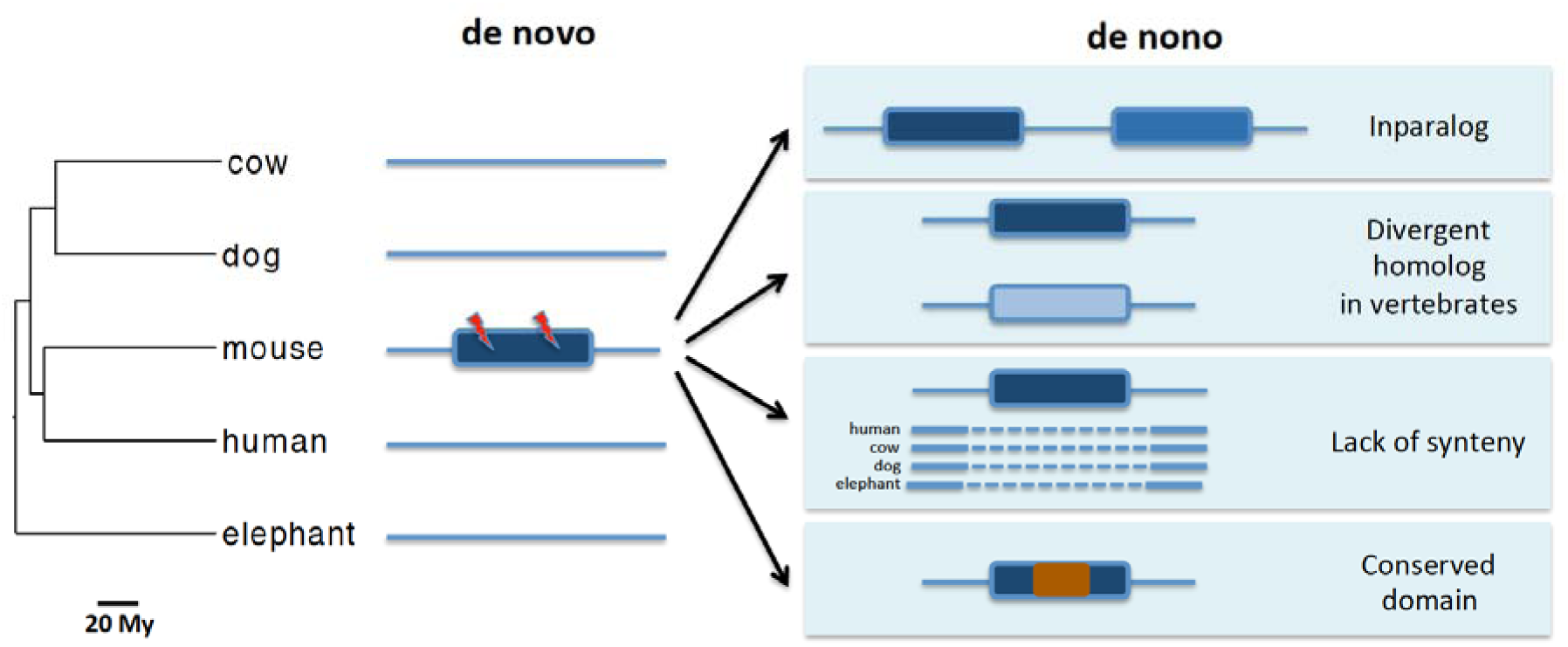
Features distinguishing de novo and ‘de nono’ genes. Rectangles, solid lines and dashed lines represent genes, nongenic syntenic regions and nonsyntenic regions, respectively. Presence of enabler substitutions (lightening bolts), absence of inparalogs and homologs in other species, conserved synteny and lack of conserved domains characterize de novo genes. Putative de novo genes that fail to conform to one or more of these criteria represent ‘de nono’ genes. My: million years.

**Figure 2.**
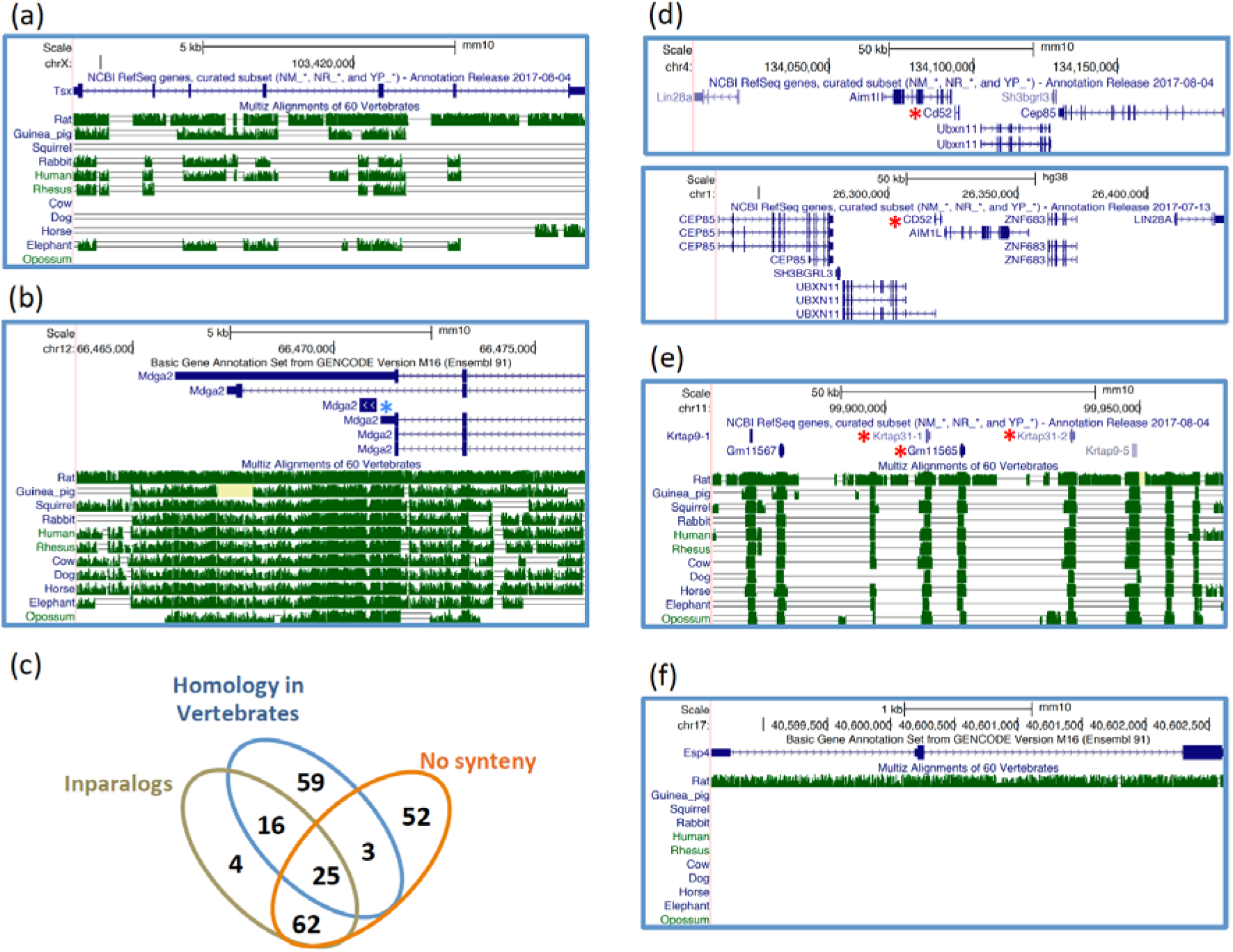
Examples of de novo and ‘de nono’ rodent genes visualized through the USCS Genome Browser. (a) An intergenic de novo gene with relatively low synteny conservation across several non-rodent mammals. (b) A de novo gene (blue asterisk) overlapping with the 3’UTR of an older gene. Notice that the gene symbol is the same for the two genes, which share no coding or protein similarity. (c) Summary of ‘de nono’ genes features. (d) A ‘de nono’ gene (*Cd52*, red asterisk) with conserved flanking genes in mouse (top) and human (bottom). (e) A tandem array of keratin-associated genes including three ‘de nono’ genes (red asterisks). (f) A ‘de nono’ gene with no synteny conservation beyond rat. Coding exons, UTRs and introns are shown as thick blue bars, thin blue bars and lines with arrows, respectively. When annotated, alternative transcripts are shown.

### Sequence similarity analyses to identify homologous genes in non-rodent genomes

According to the phylostratigraphic approach, de novo proteins in a focal taxonomic group are recognized as such because they share no significant sequence similarity with proteins from other taxa (Fig. 1). This assumption can be violated under two scenarios. First, novel proteins are routinely added to existing sequence databases, thus expanding the sequence space available to search for possible homologous sequences of PDNGs. To explore this possibility I carried out tBLASTn searches against the NCBI vertebrate nucleotide non-redundant database using mouse de novo proteins. Second, alternative sequence similarity search algorithms than those used in the original studies may reveal yet unrecognized homologs of PDNGs. For instance, profile-based approaches such as phmmer can be more accurate than non-profile methods, including BLAST, in sequence homology searches (Saripella et al. 2016). A combination of these methods has recently been applied to detect de novo genes in two yeast genomes (Vakirlis et al. 2017). I therefore interrogated a reference proteome database available through the EMBL-EBI phmmer server to identify PDNG homologs that are not recognized using BLASTP (see Methods). Finally, I visually inspected all PDNGs with synteny information to find possible orthologous in the human genome using the UCSC Genome Browser net-alignment track (Schwartz et al. 2003). Combining the results of both analyses I identified 98 PDNGs (26% of all PDNGs) with homologs in two or more vertebrate species (Table 2), including fourteen PDNGs with orthologous genes in human (Fig. 2d; Table S1).

**Table 2.**
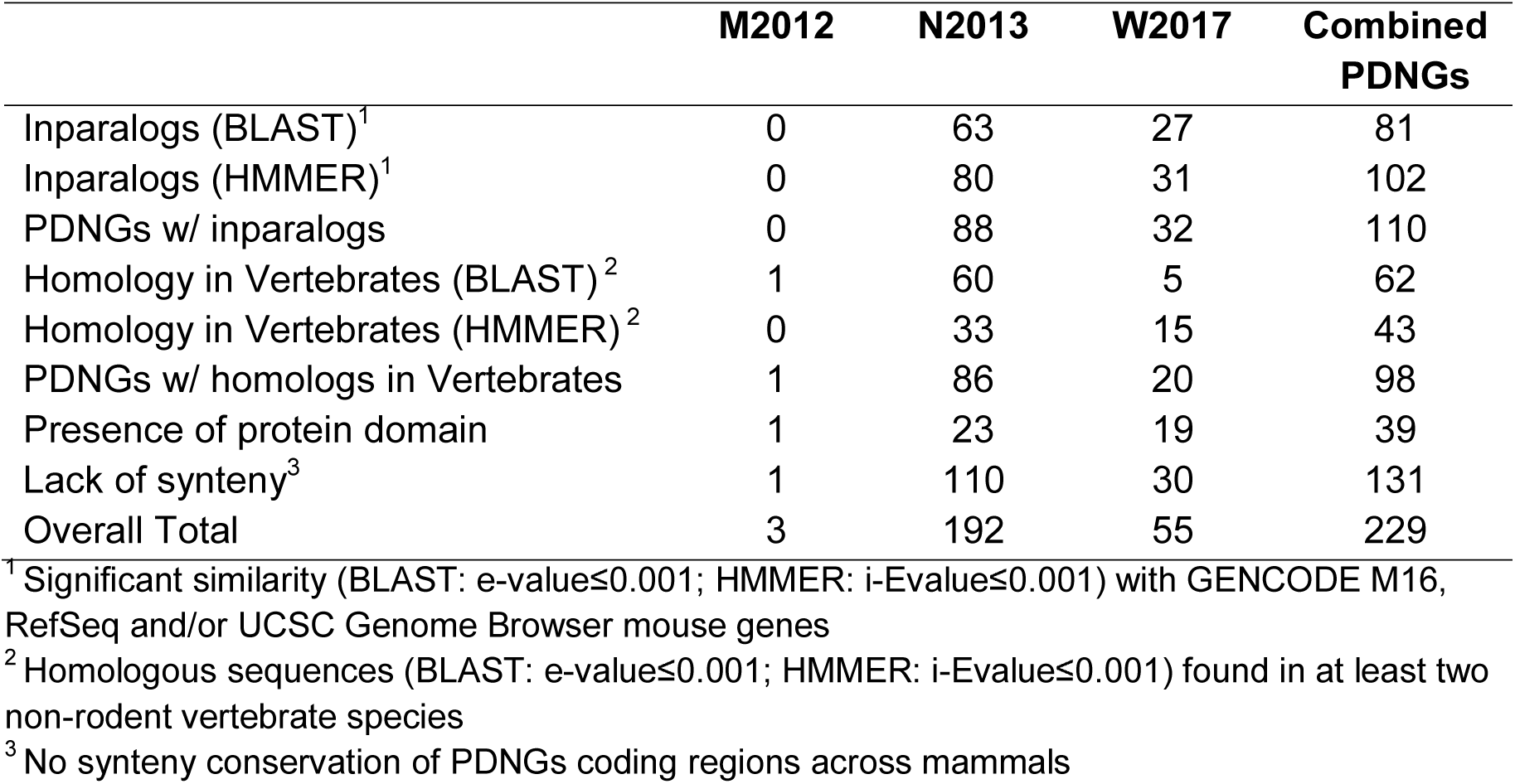
Summary of homology searches and synteny analysis for the three PDNG sets.

As expected, alignments of proteins from some of these orthologs showed short regions of sequence conservation (Fig. S1).

For five of these PDNGS with human orthologs, no inparalogs or vertebrate homologs were detected; however, genes with the same name were present in human and a visual inspection of their syntenic region, including nearby genes, confirmed that they were present in the human genome (Fig. S2).

Orthology relationships of PDNGs that were part of tandem arrays were not established given that some tandem arrays tend to experience high rates of gene turnover. Thus, the number of PDNGs with orthologs in human could be higher. In approximately one third of these PDNGs (36/107) homology to non-rodent genes was uniquely detected through the phmmer search, indicating that a significant proportion of false positives in de novo surveys will be undetected using BLAST-only approaches.

### Similarity analyses to identify paralogous genes in mouse

De novo genes can undergo duplication and form novel gene families in a genome. However, it is reasonable to consider the formation of large gene families of de novo genes unlikely in the relatively short evolutionary time period since mouse-rat divergence, which occurred as late as ∼12 million years (my) ago (Kimura et al. 2015). Perhaps more noteworthy, a conservative approach should be applied to the study of de novo genes by discarding candidates with paralogous genes in the same genome, or inparalogs. To assess the frequency of PDNGs with inparalogs, I performed similarity searches based on tBLASTn, BLASTn, and phmmer. Using the same threshold commonly applied in phylostratigraphy studies with BLAST, namely a maximum e-value of 0.001, and stringent criteria for phmmer results (see Methods), I found that 110 genes (29% of all PDNGs) have at least one paralogous gene in GENCODE M16, RefSeq, or UCSC known gene murine data sets (Fig. 2d; Table 2). Non-genic homologous sequences to PDNGs were also found using BLASTn searches against the mm10 mouse genome assembly, but were not included in the validation of PDNGs.

Fifty-four ‘de nono’ genes clustered in 20 tandem arrays, defined as groups of ‘de nono’ genes less than 100kb apart (Fig. 2e; Table S1). At least one gene pair from each array was found using a 10kb distance cutoff. Several lines of evidence suggested that these arrays were not entirely formed by de novo genes. Many arrays contained other paralogs that were not annotated as PDNGs in the first place (Fig. 2e). Moreover, PDNGs in some arrays belonged to known gene families present in other mammals. For instance, two arrays contained keratin-associated genes and another one was found within a cluster of defensin genes. Finally, most ‘de nono’ genes in arrays (37/54) showed no synteny conservation with other mammals, as expected in the presence of lineage-specific duplications rather than de novo gene formation. This finding underscores the importance of implementing more rigorous homology searches *within* the focal genome in de novo gene studies in order to remove false positives due to young gene duplicates.

### Synteny analyses

Synteny information in de novo gene investigations is crucial to detect enabler substitutions by comparing putative novel coding sequences with noncoding orthologous regions in sister taxa (McLysaght and Guerzoni 2015; Vakirlis et al. 2017). To assess synteny conservation of PDNGs, I used genome-wide alignment data available throughout the Galaxy portal. Approximately 34% PDNGs (131/381) exhibited no synteny conservation in a 60-vertebrate alignment, which includes 40 mammalian genomes (Fig. 2f; Table S1). It is arguable that some of these ‘de nono’ genes with no synteny information may represent true de novo genes that evolved in genomic region that have been lost through deletions in non-rodent mammals. However, this is unlikely given that the synteny information relies on alignments of a large number of mammalian genomes (Blankenberg et al. 2011). Given the phylogenetic distribution of the species present in the multialignment, loss of syntenic regions should have occurred independently in no less than three mammalian lineages. This also doesn’t account for possible further losses of synteny within glires (lagomorphs and rodents), which were not assessed in this study.

Many ‘de nono’ genes with no apparent orthologs outside rodents could represent lineage-specific copies of older parent genes. Indeed, 76/131 ‘de nono’ genes with no synteny conservation shared homology with at least one inparalog. Some of the remaining 55 ‘de nono’ genes with no synteny conservation may constitute rapidly evolving young gene duplicates with no detectable homology with their parent genes; for example, four of them belong to tandem arrays. As expected, the majority of ‘de nono’ genes with conserved synteny across mammals (76/98) shared homology with genes found outside rodents.

### Conserved domains in PDNGs

Conserved domains were found in protein sequences encoded by 39 PDNGs (Table S1). As expected, some of these domains belong to gene families identified in tandem arrays, such as defensing and keratin. Some conserved domains were not functionally characterized; for instance, two PDGNs encoded peptides with DUFs (domains of unknown function) and two other proteins contained a proline-rich domain. Arguably, these domains might belong to novel proteins present in multiple rodents. However, all PDNGs encoding proteins with less well-characterized domains failed to pass one or multiple other criteria to be considered valid ‘de novo’ genes (Table S1).

### New estimates of rodents de novo genes

The results of homology searches and synteny analysis showed that only 152 of the 381 PDNGs (∼40%) annotated as protein coding genes represent de novo genes (Fig. 2a-b). Notably, 70/152 (46%) of these PDNGs were automatically annotated and might represent pseudogenes (Table 1). Excluding these genes from the annotation list brings down the number of validated PDNGs to 85/314 (∼27%). Moreover, this set of 152 de novo genes is likely including several ‘de nono’ cases for three reasons. First, the criteria applied in the homology searches were particularly stringent. Non-genic paralogous sequences were excluded from inparalog searches and I required at least two non-rodent species to show significant similarity with PDNGs to identify ‘de novo’ genes. A stringent threshold was also applied in the HMMER searches (see Methods). Second, several PDNGs showed synteny with non-rodent genomes for as little as 10% of their coding regions. Third, enabler substitutions have not been searched for in the N2013 and W2017 PDNG sets (Neme and Tautz 2013; Wilson et al. 2017). Some de novo genes are also likely absent in the three studies analyzed here. Accordingly, analyses of lncRNAs in human and mouse have shown several potential protein coding de novo genes that were not been reported before (Xie et al. 2012; Chen et al. 2015; Ruiz-Orera et al. 2015; Neme and Tautz 2016).

Overall, errors in de novo gene detection depended on lack of synteny (131 genes, 34% of PDNGs), presence of mouse paralogous genes (110 genes, 29%), and/or homology with genes from at least two non-rodent vertebrates (98 genes, 26%), as summarized in Table 2. In line with results from the re-analysis of budding yeast PDNGs (Carvunis et al. 2012; Moyers and Zhang 2016), these findings call for implementing more rigorous strategies to validate PDNGs identified through phylostratigraphy using a combination of synteny analysis and extensive sequence similarity searches. Both strategies are readily implemented using existing databases and software, particularly in model taxa with established synteny data. In non-model organisms, validation of PDNGs is more problematic given the paucity of multiple closely related genomes and genome-wide alignments necessary to retrieve synteny information.

The number of confirmed de novo genes varies significantly across the three studies. Seven out of ten still annotated PDNGs from the M2012 paper were validated in this re-analysis, compared to 139/331 (42%) and 18/72 (25%) PDNGs reported in the N2013 and the W2017 studies, respectively. The two latter works were based on a phylostratigraphy-only approach, whereas Murphy and McLysaght integrated phylostratigraphy with an analysis of synteny and enabler substitutions (Murphy and McLysaght 2012). The ten annotated PDNGs all showed enabler substitutions (Murphy and McLysaght 2012). In spite of a large number of detected false positives, that is, ‘de nono’ genes, a significant difference remains in the number of validated de novo genes identified in the three studies. This is especially striking in the M2012 and N2013 studies, which focused on mouse-specific genes. Two factors seem to have contributed to the observed discrepancy between these works. First, the two analyses relied on different versions of the mouse Ensembl gene set, v56 and v66. Although both versions have been discontinued, the closest available datasets from v54 and v67 differ significantly in the number of annotated protein sequences (v54: 40,341; v67: 80,007). However, 108/117 validated de novo genes from the N2013 dataset were already present in the mouse Ensembl v54 proteome. More importantly, Murphy and McLysaght developed a pipeline incorporating several stringent filtering steps that appear to be absent in the Neme and Tautz work (Murphy and McLysaght 2012; Neme and Tautz 2013). Specifically, Murphy and McLysaght analyzed only mouse PDNGs with orthologous noncoding regions in rat and showed experimental evidence of both transcription and translation. None of these criteria were applied in the Neme and Tautz study. Therefore, the N2013 PDGNs set contains genes with weak annotation support and genes lacking synteny data. Thus, the 142 validated de novo genes from the N2013 study represent an upper boundary of the number of potential mouse de novo genes, with the caveats that some of them might be present also in the rat genome, but could not be detected in BLAST searches due to high levels of divergence. More than 3/4 of PDNGs from the W2017 study have been reclassified as ‘de nono’ genes in this study. The majority of these genes (43/54) showed significant homology with other vertebrates. Seven of them correspond to human functional orthologs and the pseudogene *Snhg11* (Fig. 2d; Fig. S2).

### Rate of de novo gene formation in mouse

In their 2013 paper, Neme and Tautz identified 780 mouse-specific putative de novo genes that, given their apparent absence in rat, would have emerged in the past ∼12 million years (Kimura et al. 2015). This corresponds to a rate of ∼65 genes/my, which is similar to mouse-specific gene duplications estimates of 63 genes/my obtained comparing mouse and Brown Norway rat (Gibbs et al. 2004) and 106 genes/my assessed using a phylogeny of 10 complete mammalian genomes (Marmoset Genome 2014). To re-calculate the rate of de novo gene formation in mouse according to my analysis, I first determined the orthology of the 152 validated de novo genes in the rat genome. Only thirteen de novo genes shared similarity to rat proteins. Thus, 139/152 validated de novo genes appear to be mouse-specific. This result implies that the maximum rate of de novo gene formation during the mouse lineage evolution correspond to ∼11.6 gene/my, assuming a mouse-rat divergence time of ∼12 million years. This indicates that de novo genes in mouse emerged at a pace that is at least about 5.4-9.1 times slower compared to gene duplicates. Notably, a recent study has shown that de novo genes originated at only ∼2.1 gene/million year in the great apes (Guerzoni and McLysaght 2016). Lower numbers of de novo genes were identified in mouse by one of these authors in the M2012 paper, suggesting that methodological differences might be largely responsible for the discrepancy in the estimates of de novo gene formation between rodents and primates.

### Characteristics of mouse de novo genes

The 152 validated de novo genes can be divided in two groups according to the quality of their annotation. The manually annotated group contained 82 de novo genes, compared to 70 automatically annotated genes (Table S2). I will refer to these two groups as MA and AA, respectively. Gene length and number of exons all increased significantly from the AA group to the MA group and for both of them in comparison to 20,391 other mouse transcripts (Table S3). These features have been observed in previous de novo gene studies, including some of the data re-examined here (Murphy and McLysaght 2012; Neme and Tautz 2013; Guerzoni and McLysaght 2016).

Seventy-one de novo genes overlapped with coding exons (20), 5’UTRs (19), 3’UTRs (9) or introns (27) of older genes (Table S4). Except for the 3’UTR cases, most de novo genes overlapped on the opposite strand of the older gene (Table S4). Given that coding exons occupy only ∼1% of mammalian genomes, the occurrence of twenty de novo genes in overlap with coding exons is especially notable. The emergence of novel ORFs on the complementary strand of preexisting genes, known as overprinting, is relatively common in viruses but is considered rare among eukaryotes (Pavesi et al. 2013). In bacteria, long ORFs tend to be present on the opposite strand of genes (Yomo and Urabe 1994). The presence of widespread long ORFs on the opposite strand of mammalian coding exons could thus accelerate the origin of de novo genes through overprinting.

Overall, de novo genes tend to overlap with other genes almost six times more often than older genes (46.7% vs. 8.4%; *P* < 0.00001, Fisher exact test). This tendency has been documented in rodents and primates (Murphy and McLysaght 2012; Neme and Tautz 2013; Ruiz-Orera et al. 2015; Guerzoni and McLysaght 2016) and implies that the evolution of de novo genes may be facilitated near older genes due to the high density of regulatory motifs, open chromatin and elevated GC content (McLysaght and Hurst 2016). The evolution of de novo genes should be facilitated in genomic regions with elevated GC content because they tend to harbor fewer AT-rich stop codons (Oliver and Marin 1996). Some of these features are also associated with de novo gene formation in intergenic regions (Vakirlis et al. 2017). Long noncoding RNAs (lncRNAs) also appear to represent another source of de novo genes, possibly because they are associated with transcriptionally active regions (Xie et al. 2012; Chen et al. 2015; Ruiz-Orera et al. 2015; Guerzoni and McLysaght 2016; Neme and Tautz 2016).

### Levels of intrinsic disorder in de novo genes and older genes

It has been argued that de novo genes encoding for proteins that show low propensity to form aggregates, and thus are less prone to induce cytotoxicity, should be more likely to be fixed (Wilson et al. 2017). Wilson and colleagues calculated the intrinsic structural disorder (ISD), a proxy for protein solubility (Monsellier and Chiti 2007; Pallares and Ventura 2016), in all mouse proteins and found that: 1) de novo genes showed the highest level of ISD, which suggested they were preadapted to become novel genes because they encode proteins with low tendency toward aggregation; 2) ISD levels increased throughout mouse genes phylostrata.

Here, I calculated ISD levels for the 152 validated rodent de novo genes and 20,391 older mouse genes using the algorithm implemented in the software PASTA (Walsh et al. 2014). Validated de novo genes showed a significantly higher proportion of ISD regions than older genes, including genes with comparable length (*P* < 0.0001, Mann-Whitney *U* test. Fig. 3a; Table S3).

**Figure 3.**
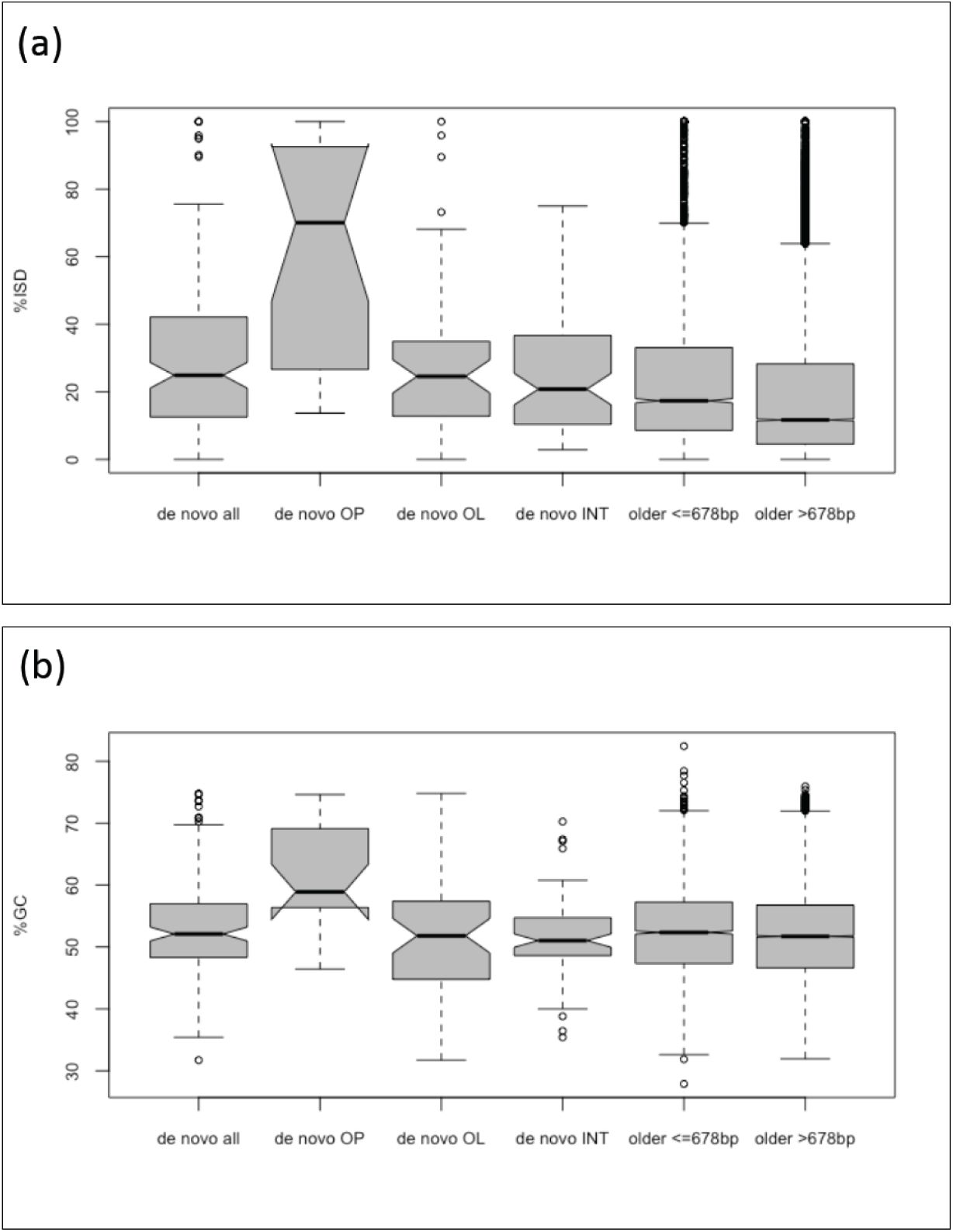
Comparison of intrinsic structural disorder percentage (a) and GC content (b) between de novo genes and older genes. Gray boxes shows values between first and third quartile. Medians are shown as black lines. Whiskers: minimum and maximum values excluding outliers. OP: overprinting. OL: overlapping with non-coding regions of older genes. INT: intergenic.

However, this derives from the particularly high levels of disorder in proteins encoded by the twenty overprinted de novo genes. ISD levels are significantly higher in these proteins compared to any other group of de novo or older proteins (all *P* < 0.002, Mann-Whitney *U* test). On the contrary, I found no significant difference in ISD levels between proteins from short older genes and proteins encoded by either intergenic or overlapping but not overprinted de novo genes (*P* = 0.051 and 0.105, respectively, Mann-Whitney *U* test).

Recent works have shown that high disorder levels in orphan and de novo proteins are associated with the elevated GC content of their genes (Basile et al. 2017; Vakirlis et al. 2017). In agreement with these findings, overprinted de novo genes showed significantly higher %GC compared to any other group of genes (all *P* < 0.0004, Mann-Whitney *U* test; Fig. 3b; Table S3), contrary to intergenic or overlapping but not overprinted de novo genes. Furthermore, the GC content and ISD levels were highly correlated in the complete set of analyzed mouse genes (*r* = 0.92, Pearson correlation). The particularly elevated GC content in overprinted de novo genes can be explained by several factors. As already mentioned, the diminished frequency of stop codons in GC-rich regions allows longer ORFs to form. Additionally, the GC content is positively correlated with the transcriptional activity in mammalian cells (Kudla et al. 2006), which could increase the likelihood of proto-genes to be spuriously expressed and eventually evolve into functional genes.

## Conclusions

The discovery of de novo genes in eukaryotes has revealed how evolutionary tinkering of noncoding regions can lead to novel protein sequences from scratch. Previous analyses relying uniquely on phylostratigraphic methods suggested that de novo genes are fixed at rates comparable to those of gene duplicates (Carvunis et al. 2012; Neme and Tautz 2013). This conclusion cannot be reconciled with the observed levels of interspecific gene homology and gene loss rates, what I referred to as the ‘gene turnover paradox’. Here, I used data from three previous studies to show that the majority of putative de novo genes thus far detected in rodents and still annotated in mouse represent either lineage-specific gene duplicates or rapidly evolving genes shared across mammals. The improved estimate of mouse-specific de novo genes points to a rate of novel gene formation that is several times lower than the gene duplication rate, a possible resolution of the ‘gene turnover paradox’. Importantly, these results also imply that the known homology detection bias in phylostratigraphy is *not* minimized by focusing on the youngest genes in a given species, as previously suggested (Wilson et al. 2017). However, as shown in this and other studies (Murphy and McLysaght 2012; Vakirlis et al. 2017), false positive rates in de novo gene surveys can be significantly reduced by utilizing a combination of more sensitive homology search approaches and synteny analyses.

In one of the re-examined studies, Wilson and co-authors (2017) found that putative de novo proteins have the highest levels of intrinsic structural disorder (ISD), a measure that negatively correlates with protein toxicity, among mouse proteins. This would suggest that de novo genes evolve more frequently from proto-genes that are preadapted because they encode peptides with low level of toxicity. Mouse de novo proteins validated in my study also show higher ISD levels than older genes; however, I found that this is due to a subset of de novo genes that share high GC content and overlap with coding exons of older genes. In agreement with recent observations (Basile et al. 2017; Vakirlis et al. 2017), this shows that the elevated disorder of mouse de novo proteins represent a mere consequence of the high %GC of some de novo genes, rather than supporting the preadaptation hypothesis.

## Methods

*Putative* de novo *genes*

### Murphy and McLysaght 2012

Mouse de novo gene IDs were retrieved from Table 1 of the Murphy and McLysaght study (Murphy and McLysaght 2012). These genes were found using the Ensembl version 56. Protein-coding genes from the two closest available Ensembl versions, v54 and v67, were downloaded from the Ensembl archives (https://www.ensembl.org/info/website/archives/index.html). Out of the 69 putative mouse de novo genes, only twenty-six were still annotated as protein-coding genes in v67.

### Neme and Tautz 2013

Neme and Tautz identified de novo gene using the mouse Ensembl version 66. We retrieved all the 80,007 mouse transcript and protein IDs and sequences annotated in the closest available data set, the archived Ensembl version 67. We found a match for 779 out of 780 mouse putative de novo genes in the Ensembl v67 version and selected the longest protein isoform for these genes for subsequent analyses. Six PDNGs were removed from the 779 gene set after applying a minimum protein length threshold of 30 amino acids leading to a total of 773 analyzed PDNGs in the N2013 data set. The presence of N2013 PDNGs in the mouse Ensembl v54 proteome was determined using a tBLASTn search with an evalue threshold of 0.001. Hits that shared at least 90% sequence identity over at least half of the query were considered orthologous sequences.

### Wilson at al. 2017

Ensembl gene IDs and sequences of the 84 mouse young genes were obtained from the supplementary table 2 of the Wilson et al. paper (Wilson et al. 2017). Transcript and protein IDs and sequences annotated in the closest available data set, the archived Ensembl version 75 (https://www.ensembl.org/info/website/archives/index.html),the same used in the W2017 paper.

#### Updated annotation of PDNGs

The UCSC Genome Browser and Table Browser have been used to retrieve genome coordinates of PDNGs from the three papers’ datasets (http://genome.ucsc.edu/cgi-bin/hgTables). However, the Ensembl gene track from the most recent mouse genome assembly (GRCm38/mm10, December 2011) does not contain all Ensembl IDs corresponding to PDNGs. Genome coordinates of PDNGs were therefore retrieved using the previous mouse genome assembly (NCBI37/mm9, July 2007). These coordinates were then transformed into coordinates of the mouse mm10 assembly using the LiftOver tool (http://genome.ucsc.edu/cgi-bin/hgLiftOver). The genome coordinates of most PDNG coding exons were successfully lifted to the mm10 assembly and uploaded to the Galaxy portal (https://usegalaxy.org). Genome coordinates of the coding exons of mouse RefSeq, GENCODE M16 (same gene set as Ensembl v91) and UCSC ‘known’ genes were also uploaded to Galaxy. PDNG coding exons were then joined using the Galaxy tool ‘Join’ in the Menu ‘Operate on Genomic Intervals’ applying ‘All records of first dataset’ to return their overlap with coding exons of each of the three gene sets with a minimum of 50 bp overlap (all PDNGs had at least one coding exon longer than 50bp). Coding exons of 381 PDNGs overlapped with coding exons of at least one gene from the three used gene sets. Notice that overlapping genes on opposite strands of PDNGs were not excluded. PDNG whose IDs were not available in the mm9 Ensembl track were re-annotated by querying their coding sequences against the mm10 genome assembly using blat (http://genome.ucsc.edu/cgi-bin/hgBlat) to visually find matches with annotated GENCODE M16, RefSeq and UCSC known genes. The M2012 and W2017 PDNGs were visually inspected on the UCSC Genome Browser for overlap with annotated genes. Annotation was confirmed for 9/69 M2012 PDNGs (Table 1).

#### Sequence similarity analyses to identify paralogs

Several types of BLAST searches on multiple mouse databases were carried out using a consistent e-value threshold of 0.001. The mouse genome assembly mm10 was searched locally using tBLASTn (Camacho et al. 2009). In searches against the mouse genome only hits against the coding sequence of known genes were considered valid paralogous genes of putative de novo genes. The combined mouse proteomes from GENCODE M16 genes (Harrow et al. 2012), the RefSeq genes (O’Leary et al. 2016) and the UCSC Genome Browser ‘known genes’ (http://genome.ucsc.edu/cgi-bin/hgTables) databases were searched locally using BLASTP. BLAST results were parsed and filtered using perl scripts and Unix commands. Matches over less than 50% of the query sequence were removed to increase stringency. Matches of PDNGs with multiple proteins were carefully inspected to ensure that they corresponded to multiple loci rather than alternative transcripts of the same gene. Alignments of PDNGs with a single match were also inspected to determine if these hits represented paralogous genes rather than self-hits.

Similar searches were performed locally using the algorithm phmmer in the HMMER suite (http://hmmer.org/). Each PDNG protein set was queried against the three combined GENCODE, RefSeq and USCS proteomes using default parameters. Results were visually inspected to identify matches between protein sets. Hits with c-Evalue and i-Evalue below 0.001 were considered positive matches.

#### Sequence similarity analyses to identify homologs in vertebrates

The vertebrate (taxid:7742) NCBI nr protein database was interrogated between September 2017 and January 2018 in the NCBI BLAST portal (https://blast.ncbi.nlm.nih.gov/Blast.cgi) using default settings except an e-value threshold of 0.001, excluding Rodents (taxid:9989). The reference proteomes database (https://www.ebi.ac.uk/reference_proteomes) was interrogated between September 2017 and January 2018 using phmmer with a higher than default stringency e-value of 1e^-05^ and excluding rodents from the search (https://www.ebi.ac.uk/Tools/hmmer/search/phmmer). Only PDNG proteins with significant hits with proteins from at least two vertebrates were considered positive matches in both BLAST and HMMER searches.

#### Synteny analyses

Mouse genome coordinates in BED format of the PDNGs coding exon were retrieved from the UCSC Genome Browser table browser tool (http://genome.ucsc.edu/cgi-bin/hgTables) using either Ensembl identifiers or novel RefSeq/UCSC identifiers from the re-annotation of the three datasets (Tables S1-2). The BED coordinates were then sent to the Galaxy portal (https://usegalaxy.org).

I generated a workflow on Galaxy (https://usegalaxy.org/u/claudiocasola/w/maf-blocks-for-mouse-mm10-sequences) to obtain MAF blocks (Multiple Alignment Format blocks) from aligned sequences in the mouse genome assembly mm10 (Blankenberg et al. 2011). Briefly, the workflow utilizes genome coordinates to extract MAF blocks from the 100-way multiZ alignment based on the human genome assembly hg19. Overlapping MAF blocks were merged, filtered to retain only mouse blocks, and joined to the coordinates of each coding exon of the putative de novo genes. A few remaining overlapping MAF blocks were manually removed from the MAF datasets.

#### Protein domain analyses

The NCBI Conserved Domain repository (Marchler-Bauer et al. 2017) was interrogated with proteins encoded by PDNGs between 09-2017 and 01-2018 using default parameters except inclusion of retired sequences in the batch search portal (https://www.ncbi.nlm.nih.gov/Structure/bwrpsb/bwrpsb.cgi).

Conserved domains of PDNGs with no evidence paralogy and lack of synteny were also searched throughout the InterPro server (https://www.ebi.ac.uk/interpro/) in March 2018.

#### Gene structure

Gene length, coding length, intron length, UTRs length and exon number of mouse genes were obtained from the GENCODE M16 dataset through the UCSC Table Browser. The 64,506 transcripts were filtered to remove noncoding sequences and genes with only automatic annotation. Transcripts matching PDNGs were also removed. All except the shortest transcripts of the remaining genes were removed, leaving 20,470 genes.

#### Quality of gene annotation

The annotation quality of validated de novo genes was assessed by retrieving data from the GENCODE M16 Basic gene set and the UCSC known genes from the UCSC Table Browser. GENCODE transcript support levels range from 1 (all splice junctions of the transcript are supported by at least one non-suspect mRNA) to NA (the transcript was not analyzed). Additionally, high-quality GENCODE genes are manually annotated in HAVANA (https://www.gencodegenes.org/gencodeformat.html). Fourty-two de novo genes with either no HAVANA ID or transcript support level equal NA were considered low-quality genes. Similarly, 31 UCSC known transcript that has not been reviewed or validated by the RefSeq, SwissProt or CCDS staff were considered low quality.

#### Protein disorder and protein aggregation analyses

The software PASTA 2.0 (Walsh et al. 2014) with default settings was used to estimate intrinsic structural disorder (ISD) in proteins encoded by the three PDNG sets and 20,391 non-de novo Ensembl v91 proteins.

#### Estimates of gene duplication rates

Gene duplication events estimated to have occurred in mouse since its divergence from the Brown Norway rat have been obtained from Worley et al. (2014). Gene duplications and losses were modeled using the maximum-likelihood framework implemented in the CAFE package (Han et al. 2013). A total of 1,052 mouse-specific gene duplications were calculated based on gene family data totaling 18,215 genes (supplementary figure 7 in (Marmoset Genome 2014)). Assuming ∼22,000 genes in the mouse genome, the actual overall number of gene duplicates is 1,275, leading to a rate of duplication of ∼106/my.

#### Overlap between genes

Each validated de novo genes was visually inspected through the UCSC Genome Browser to identify possible overlap with other genes. To find the genome-wide proportion of overlapping genes, transcripts from all genes in the mouse GENCODE M16 basic gene set were downloaded from the UCSC Table Browser. Transcripts from the mitochondrial genome, non-protein coding transcripts and transcripts from de novo genes were removed. For each remaining gene, only the longer transcript was retained, leaving a total of 22,396, of which 1876 (∼8.4%) overlapped other genes.

#### Rat de novo gene orthologs

The genome coordinates of the 152 validated de novo genes from the mouse assembly mm10 were used to retrieve syntenic regions in the rat rn6 assembly with the LiftOver tool in the UCSC Genome Browser. A total of 29,107 Ensembl and transcript coding regions were downloaded using the UCSC Table Browser (Data last updated: 2017-06-09). Proteins and CDS were searched against the retrieved rat genomic regions syntenic with mouse de novo genes using tBLASTn, e-value threshold = 0.001. The BLAST results were parsed using an in-house perl script and filtered to retain hits longer than 30bp and with at least 97% DNA sequence identity between CDS and genome. This step left 67 putative orthologous proteins to mouse de novo genes. Some of these proteins were orthologous to proteins that in mouse overlap to the validated de novo genes. Thus, I manually inspected these proteins against the mouse mm10 assembly using BLAT in the UCSC Genome Browser and by running a BLASTP search between the 152 mouse de novo proteins and the 67 candidate rat orthologs. The same approach was used to retrieve 17,619 rat RefSeq proteins and CDS. I obtained 168 candidates that were screened against the 133 mouse validated de novo genes. Additionally, I searched the combined 235 candidate Ensembl and RefSeq proteins against the 152 mouse de novo proteins using phmmer locally with default settings.

## Acknowledgments

I am grateful to Aaron Quinlan and Ryan Layer for allowing me to use the term ‘de nono’. I thank Michelle Lawing for help with statistical analyses and for comments on the manuscript. This work has been supported by the National Institute of Food and Agriculture, U.S. Department of Agriculture, under award number TEX0-1-9599, the Texas A&M AgriLife Research, and the Texas A&M Forest Service.

