## Supplementary Materials for "From de novo to ‘de nono’: most novel protein coding genes identified with phylostratigraphy represent old genes or recent duplicates"

**Claudio Casola^1*^**

**^1^** Department of Ecosystem Science and Management

Texas A&M University, College Station, TX 77843-2138

* Corresponding author

495 Horticulture Rd, College Station, 77843-2138 TX

(979) 845-8803

**Supplementary Table 3. Gene features of *bona fide* de novo genes and other genes.**

|  | # | Gene Length | CDS Length | # Exons | %ISD | %GC |
| --- | --- | --- | --- | --- | --- | --- |
| Validated de novo | 155 | 9,375 | 406 | 2.5 | 32.5 | 52.8 |
| MA de novo | 85 | 7,835 | 413 | 2.9 | 29.5 | 51.3 |
| AA de novo | 70 | 11,180 | 398 | 2.0 | 36.1 | 54.6 |
| Overlapping | 74 | 6,592 | 399 | 2.3 | 37.3 | 54.5 |
| Intergenic | 81 | 11,815 | 413 | 2.7 | 28.0 | 51.3 |
| Overlapping non-CDS | 51 | 5,924 | 382 | 2.1 | 29.8 | 52.0 |
| Overprinting (CDS) | 20 | 8,295 | 441 | 2.9 | 57.7 | 61.2 |
| Others ≤678 bp | 3,806 | 15,222 | 475 | 4.1 | 24.4 | 52.3 |
| Others all | 16,585 | 52,495 | 1934 | 11.2 | 19.1 | 51.7 |

Averages are shown for each parameter. %ISD: percentage of intrinsic structural disorder.

**Supplementary Table 4. Overlap of *bona fide* de novo genes with older genes.**

|  | **CDS** | **5’UTR** | **3’UTR** | **Intron** | **Total** |
| --- | --- | --- | --- | --- | --- |
| MA opposite | 10 | 4 | 1 | 12 | 41 |
| MA same | 0 | 1 | 7 | 3 | 8 |
| AA opposite | 10 | 14 | 1 | 11 | 19 |
| AA same | 0 | 0 | 0 | 1 | 1 |
| TOTAL opposite | 20 | 18 | 2 | 23 | 59 |
| TOTAL same | 0 | 1 | 7 | 4 | 12 |
| **TOTAL all** | **20** | **19** | **9** | **27** | **75** |

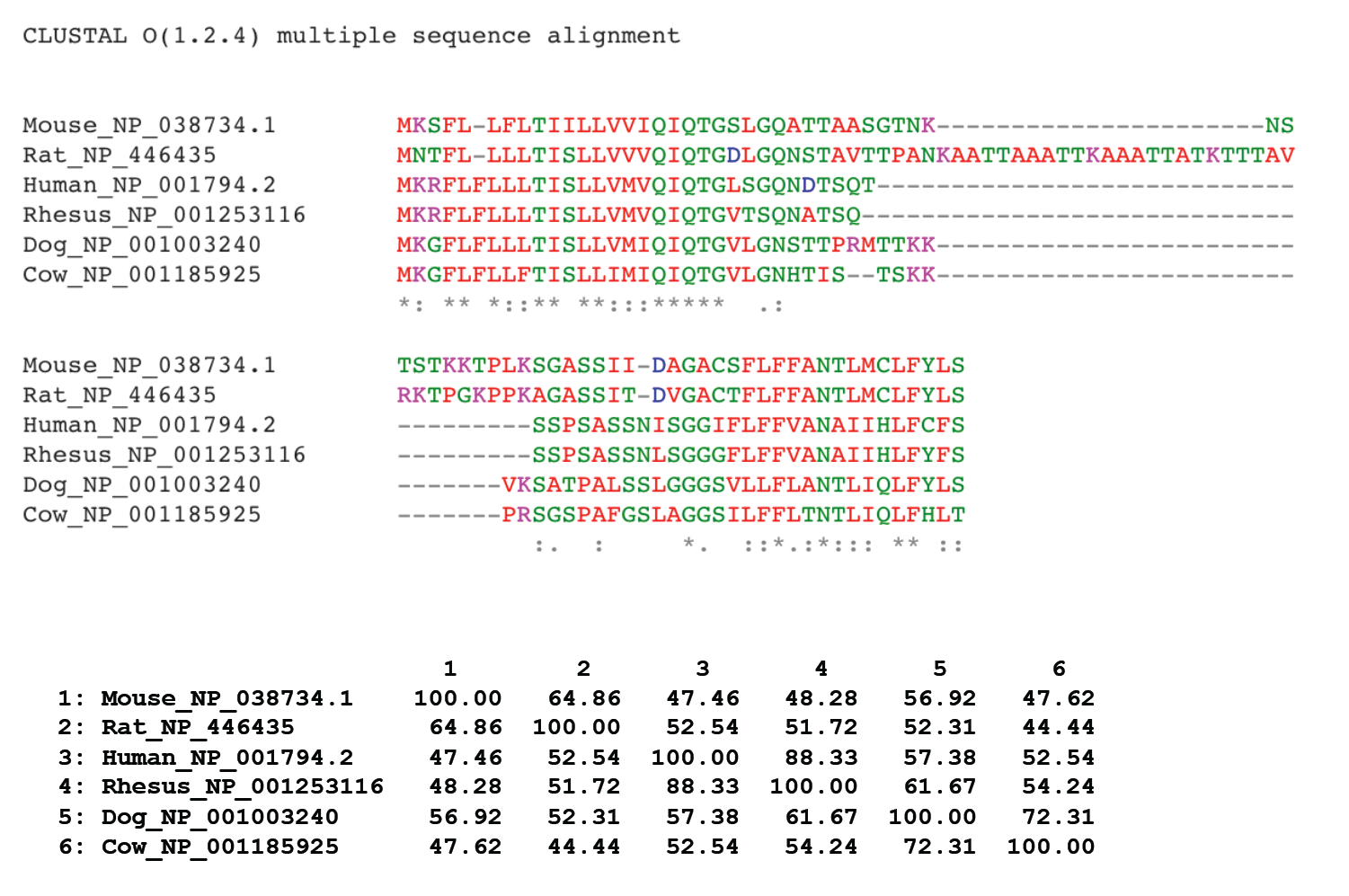

**Supplementary Figure 1.** Protein alignment of Cd52 genes from several mammals. The table shows pairwise percentage identities of the protein sequences.

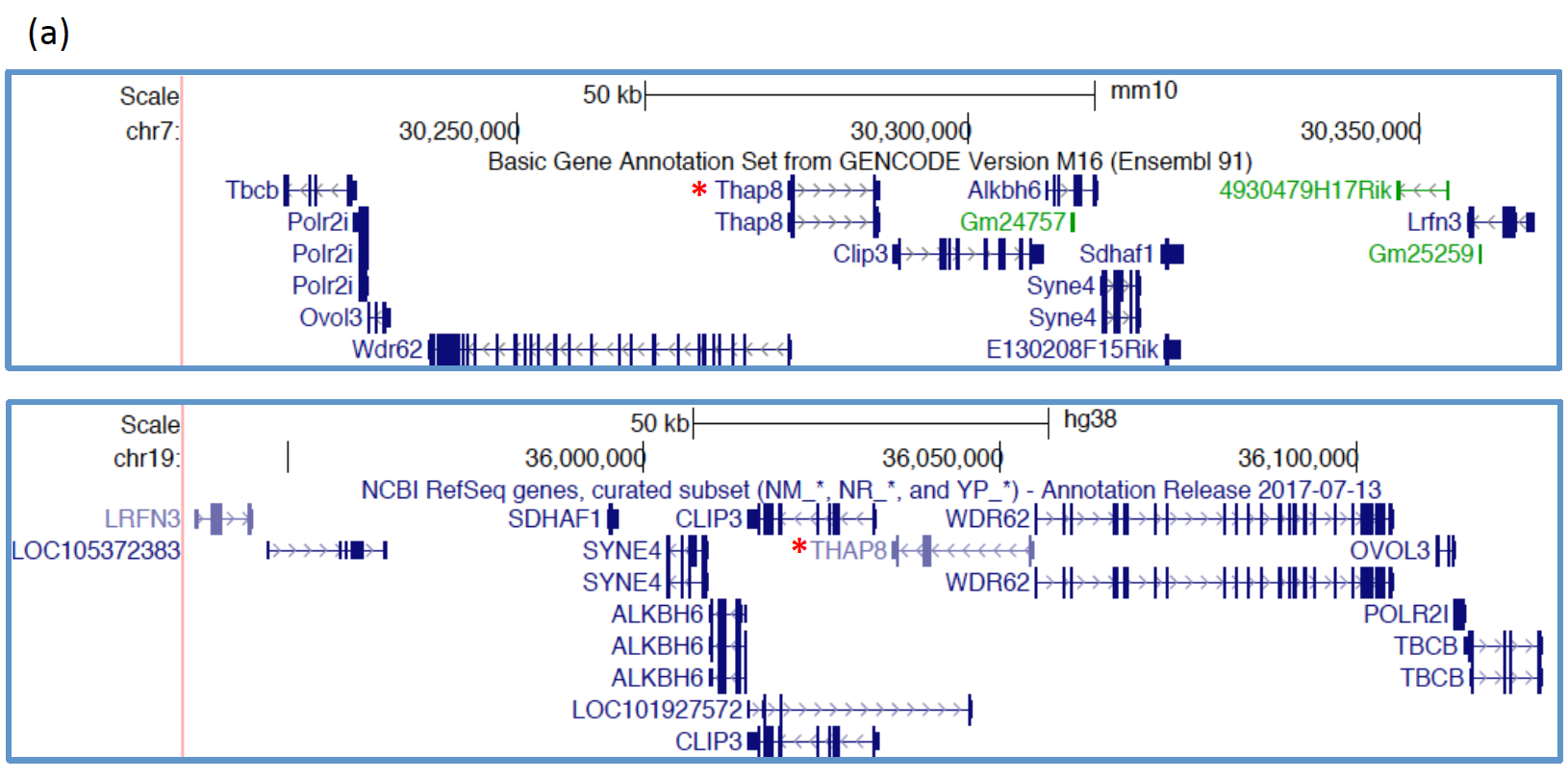

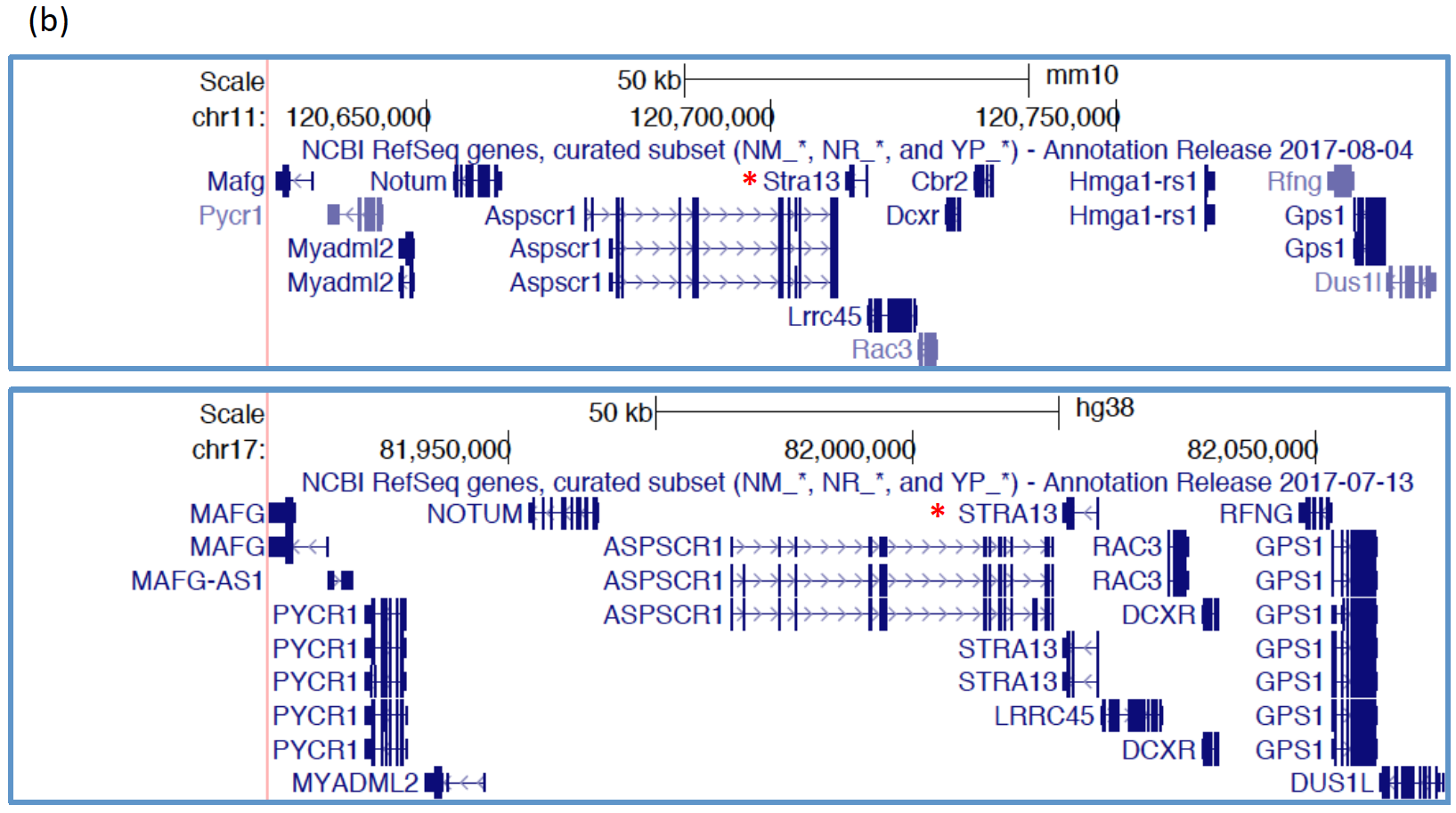

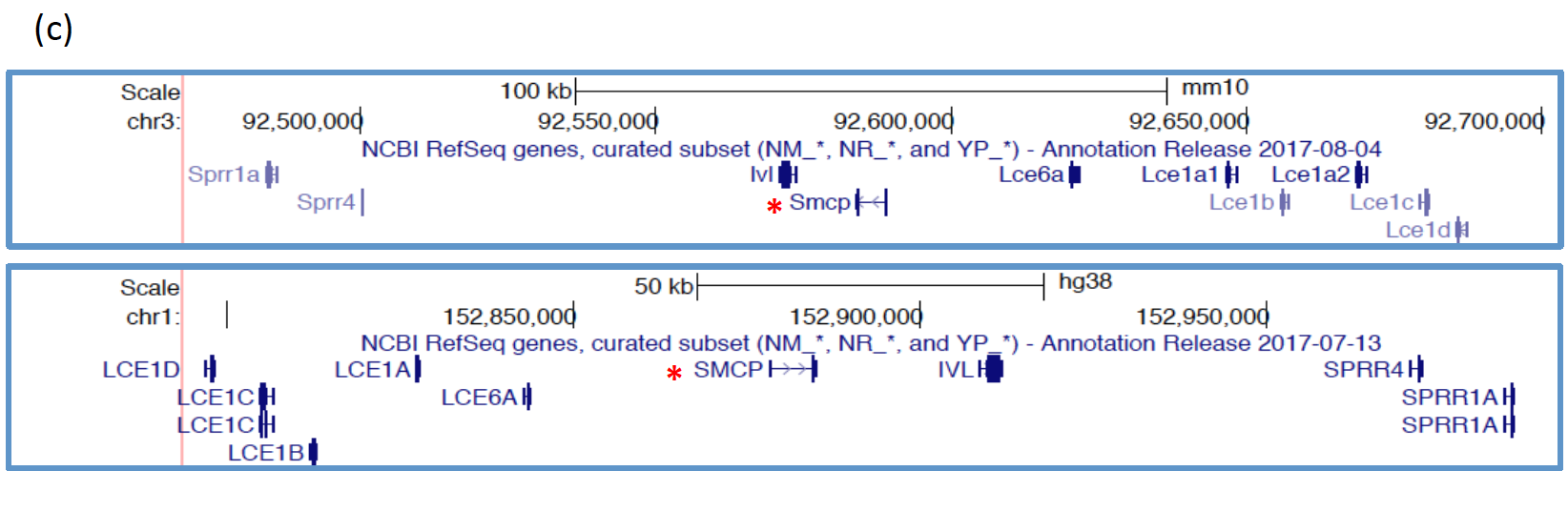

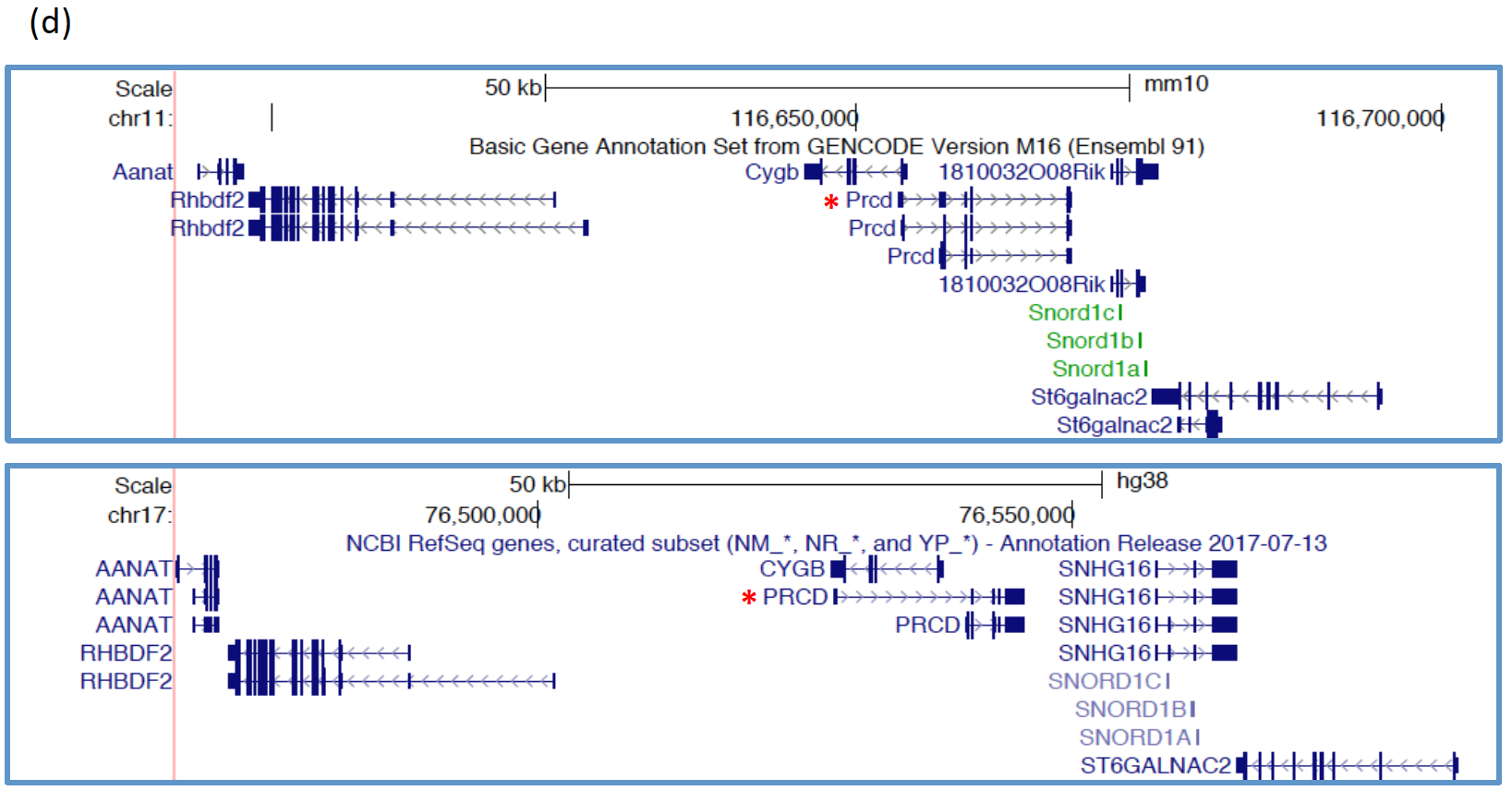

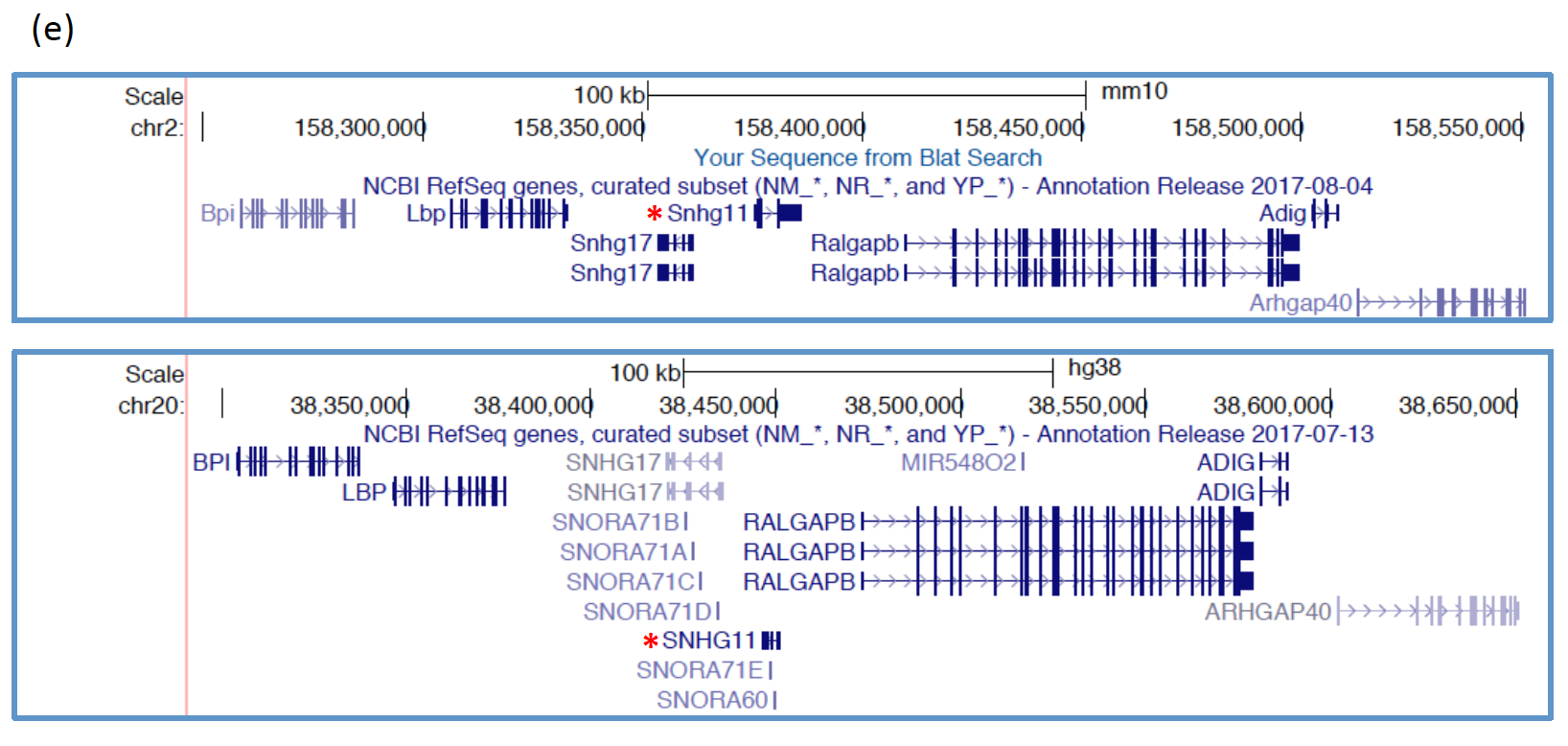

**Supplementary Figure 2.** Synteny conservation between mouse and human genomic regions surrounding five de nono genes with no detectable homology within the mouse genome and among non-rodent vertebrates. Top figure: mouse genome assembly mm10. Bottom figure: human genome assembly hg13. (a) Thap8; (b) Stra13; (c) Smcp; (d) Prcd; (e) Snhg11. Red asterisks are shown next to de nono genes. Coding exons, UTRs and introns are shown as thick blue bars, thin blue bars and lines with arrows, respectively. When annotated, alternative transcripts are shown.
